# Regulation of ATM and ATR by Smarcal1 and BRG1

**DOI:** 10.1101/261610

**Authors:** Ramesh Sethy, Radhakrishnan Rakesh, Ketki Patne, Vijendra Arya, Tapan Sharma, Dominic Thangminlen Haokip, Reshma Kumari, Rohini Muthuswami

## Abstract

The G2/M checkpoint is activated on DNA damage by the ATM and ATR kinases that are regulated by post-translational modifications. In this paper, the transcriptional co-regulation of *ATM* and *ATR* by SMARCAL1 and BRG1, both members of the ATP-dependent chromatin remodeling protein family, is described. SMARCAL1 and BRG1 co-localize on the promoters of *ATM* and *ATR*; downregulation of SMARCAL1/BRG1 results in transcriptional repression of *ATM/ATR* and therefore, overriding of the G2/M checkpoint leading to mitotic abnormalities. On doxorubicin-induced DNA damage, SMARCAL1 and BRG1 are upregulated and in turn, upregulate the expression of ATM/ATR.

Phosphorylation of ATM/ATR is needed for the transcriptional upregulation of *SMARCAL1* and *BRG1*, and therefore, of *ATM* and *ATR* on DNA damage. The regulation of ATM/ATR is rendered non-functional if SMARCAL1 and/or BRG1 are absent or if the two proteins are mutated such that they are unable to hydrolyze ATP, as in for example in Schimke Immuno-Osseous Dysplasia and Coffin-Siris Syndrome. Thus, an intricate transcriptional regulation of DNA damage response genes mediated by SMARCAL1 and BRG1 is present in mammalian cells.

## INTRODUCTION

ATM and ATR, both members of the PI3K family, are key regulators of DNA damage response pathway [1,2]. ATM is recruited to the site of double-strand breaks by the MRN complex [3] while ATR is recruited to the site of single-strand breaks by ATRIP and RPA [4,5]. Both proteins then proceed to phosphorylate multiple target proteins effecting cell cycle arrest and activating the DNA damage response pathway [6,7]. Despite overlapping substrates, it is known that only ATR is essential for the viability of replicating cells [8]. ATM, many experiments have shown, is nonessential [9].

ATM and ATR are regulated by phosphorylation. Activation of ATM kinase is dependent on the MRN complex as the protein directly interacts with the complex and this interaction is needed for the activation of the kinase [3,10]. ATM is present as an inactive homodimer in the cells; autophosphorylation on serine 1981 leads to the formation of the active monomers [11]. ATM autophosphorylation is regulated by the phosphatase, PP2A, as overexpression of dominant negative mutant of PP2A results in constitutive autophosphorylation on serine 1981 even in the absence of DNA damage [12,13]. Another phosphatase, WIP1, has also been shown to regulate the activity of ATM [12,14]. Recently, Tip60 complex has been shown to acetylate ATM in response to DNA damage, thus, adding one more post-translational modification to the regulatory repertoire [15,16].

ATM can recruit ATR to the sites of DNA damage [17,18]. ATR too is regulated by phosphorylation; however, the kinase and the phosphatase acting on ATR in the DNA damage response pathway have not been completely delineated.

The DNA damage response pathway is also controlled by the ATP-dependent chromatin remodeling proteins [19–21]. In particular, BRG1, a known transcription modulator, has been shown to participate in the DNA damage response pathway [22–24]. Mutations in BRG1 is associated both with cancer and Coffin-Siris Syndrome (CSS) [25–27]. SMARCAL1, another member of the family, in complex with RPA, has also been shown to stabilize stalled replication fork when DNA is damaged [28–33]. Studies have also shown that ATR phosphorylates SMARCAL1 and thus, regulates its activity [34]. Mutations in SMARCAL1 has been linked to Schimke Immuno-osseous Dysplasia (SIOD) [35]. However, the role of SMARCAL1 and BRG1 in regulating ATM and ATR kinases has not been studied as yet.

The results presented in this paper originated from the observation that downregulation of either *SMARCAL1* or *BRG1* in HeLa cells resulted in the formation of mitotic cells possessing multipolar spindles. Further, both *SMARCAL1* and *BRG1* downregulated cells showed DNA damage even in the absence of any DNA damage-inducing agent. Knowing that SMARCAL1 and BRG1 mutually co-regulate the expression of each other [36] and that these proteins function as transcriptional co-regulators, the expression of cell cycle checkpoint genes was investigated. In this paper, the transcriptional co-regulation of *ATM* and *ATR* by SMARCAL1 and BRG1 is reported. Downregulation of either *SMARCAL1* or *BRG1* results in downregulation of *ATM* and *ATR* leading to the failure of the G2/M checkpoint. During DNA damage, the expression of *SMARCAL1* and *BRG1* is upregulated. This upregulation results in increased expression of *ATM* and *ATR*, thus, creating a transcriptional network that governs the DNA damage response pathway. The transcriptional network is feedback regulated by the phosphorylation status of ATM and ATR. Finally, it is shown that the SIOD-associated mutations in SMARCAL1 and CSS-associated mutations in BRG1 result in abrogation of the DNA damage response pathway due to their inability to synthesize ncRNA when DNA damage is induced.

## RESULTS

### Stable downregulation of *SMARCAL1* and *BRG1* in HeLa cells results in mitotic abnormalities

*SMARCAL1* and *BRG1* are a pair of mutually co-regulated genes in HeLa cells that encode for SMARCAL1 and BRG1, members of the ATP-dependent chromatin remodeling family [36]. To understand the impact of this co-regulation on the cell, *SMARCAL1* and *BRG1* were stably downregulated in HeLa cells using shRNA construct against the genes. Both Sh*SMARCAL1* as well as Sh*BRG1* cells exhibited multipolar spindles and multinucleate cells (Fig. 1A-D and Supplementary Fig. 1A and B). However, these stably downregulated cells could not be stored for a long time. Therefore, all the experiments described, henceforth, in this paper were done using transient transfections.

**Figure 1.**
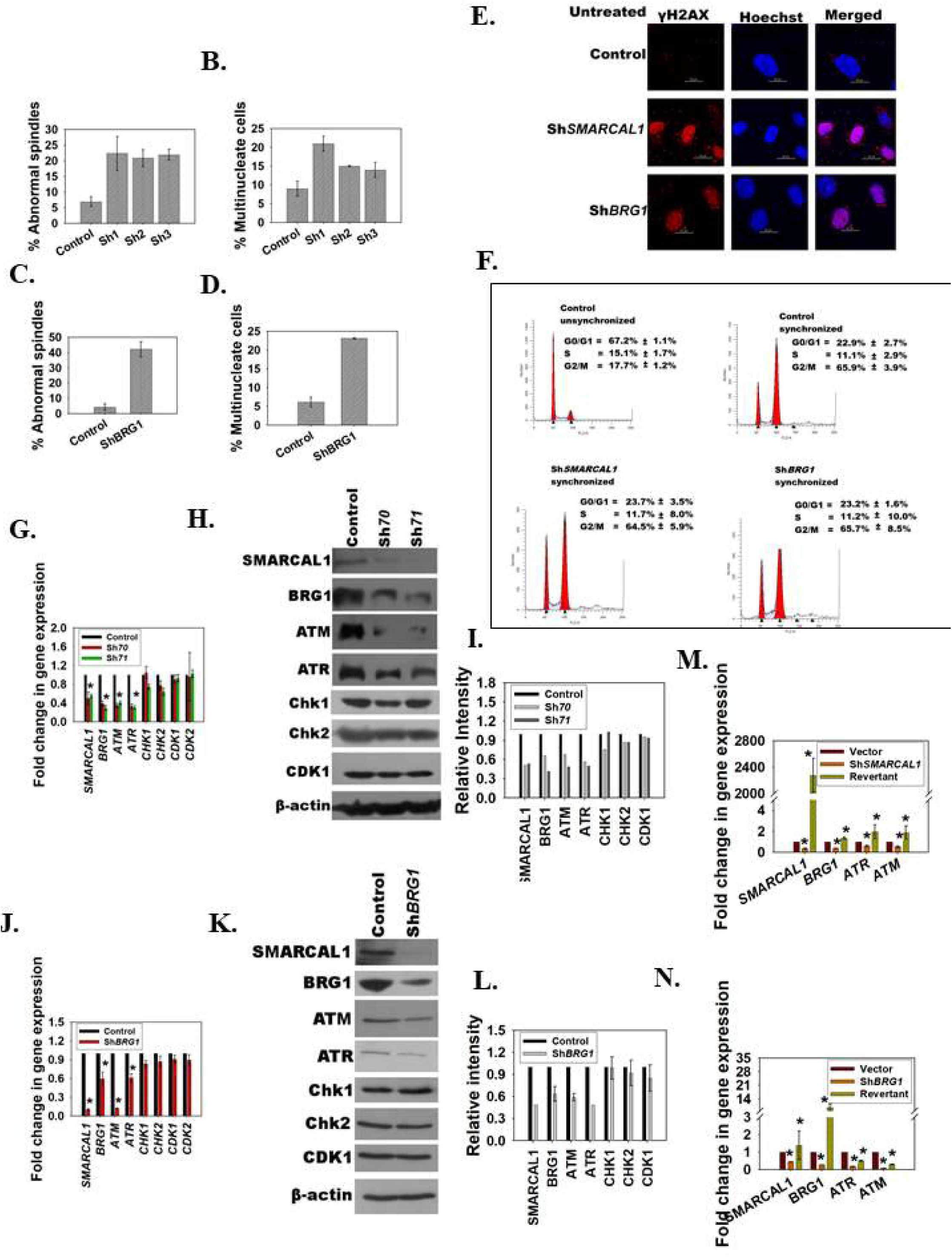
*SMARCAL1* and *BRG1* transcriptionally regulate *ATM* and *ATR*. (A). Abnormal spindle formation was quantitated in control and *SMARCAL1* downregulated cells. (B). Multinucleate cells were quantitated in control and *SMARCAL1* downregulated cells. (C).Abnormal spindle formation was quantitated in control and Sh*BRG1* cells. (D). Multinucleate cells were quantitated in control and Sh*BRG1*cells. In these experiments, stable transfectants were used. In case of *SMARCAL1*, three clones (Sh1, Sh2, and Sh3) were analysed while in case of *BRG1*, one clone was analysed. The data is presented as an average ±s.d. of two independent experiments where >100 cells were counted in each experiment. (E). The formation of γH2AX foci was analysed in control, Sh*SMARCAL1*, and Sh*BRG1* cells in the absence of DNA damaging agent. In this experiment, transiently transfected cells were analysed. (F). The progression of control, SMARCAL1 downregulated cells (Sh *SMARCAL1)* and BRG1 downregulated cells (Sh *BRG1)* through the cell cycle was monitored using FACS. Transiently transfected cells were synchronized using double thymidine block as explained in materials and methods. (G). The expression of genes regulating the G2/M checkpoint was analysed by qPCR in *SMARCAL1* downregulated cells. (H). The expression of proteins regulating the G2/M checkpoint were analysed by western blot in SMARCAL1 downregulated cells. (I). Quantitation of the western blots. (J). The expression of genes regulating the G2/M checkpoint was analysed by qPCR in Sh*BRG1* cells. (K). The expression of proteins regulating the G2/M checkpoint were analysed by western blot in Sh*BRG1* cells. (L). Quantitation of the western blots. (M). The expression of *SMARCAL1*, *BRG1*, *ATM* and *ATR* was analysed after transfecting cells either with Sh *SMARCAL1* (targeting the 3’UTR of the gene) or co-transfecting cells with Sh *SMARCAL1* (targeting the 3’UTR of the gene) along with *SMARCAL1* overexpression cassette. (N). The expression of *SMARCAL1*, *BRG1*, *ATM* and *ATR* was analysed after transfecting cells either with Sh*BRG1* (targeting the 3’UTR of the gene) or co-transfecting cells with Sh*BRG1* (targeting the 3’UTR of the gene) along with *BRG1* overexpression cassette. The data in case of qPCR experiments is presented as average ±s.d of three independent experiments (*P < 0.001; unpaired student’s t-test).

Downregulation of SMARCAL1 has been reported to increase replication stress [29] and indeed, increased γH2AX foci was found even in the absence of any DNA damaging agent in both Sh*SMARCAL1* and Sh*BRG1* cells as compared to the control cells indicating endogenous DNA damage in these downregulated cells (Fig. 1E). Increased DNA damage should result in G2/M arrest due to activation of G2/M checkpoint [42]. However, Sh*SMARCAL1* and Sh*BRG1* cells did not arrest at the G2/M boundary; instead, FACS analysis showed that the population of G0/G1, S, G2/M cells were similar in the control, Sh*SMARCAL1*, and Sh*BRG1* cells, leading us to hypothesize that G2/M checkpoint was possibly not functional when SMARCAL1/BRG1 was downregulated in HeLa cells (Fig. 1F).

### ATM and ATR expression is downregulated in both Sh*SMARCAL1* and Sh*BRG1* cells

The G2/M checkpoint is mediated by ATM and ATR kinases [43,44] in response to DNA damage. On activation these kinases phosphorylate Chk1 and Chk2 that subsequently phosphorylate Cdc25 and blocks the activation of cyclinB-Cdk1 complex [45]. As both SMARCAL1 and BRG1 are transcription factors, it was hypothesized that the expression of some of the genes encoding for proteins involved in the G2/M checkpoint was altered resulting in the inactivation of the checkpoint. Hence, the transcription profile of genes encoding for G2/M checkpoint were estimated in cells transiently transfected with shRNA either against *SMARCAL1* (Sh*70* and Sh*71*) or against *BRG1* (Sh*BRG1*). In case of *SMARCAL1*, two independent shRNA, namely Sh*70* and Sh*71*, were used to ensure the observed results were not due to off-target effect. Both qPCR and western blots showed that the expression of ATM and ATR were downregulated in Sh*SMARCAL1* as well as Sh*BRG1*cells (Fig. 1G-L). However, the levels of *Chk1*, *Chk2*, *CDK1* and *CDK2* were found to be unchanged (Fig. 1G-L). To confirm that the regulation of *ATM* and *ATR* was due to *SMARCAL1* and *BRG1* only, ShRNA constructs targeting the 3’ UTR of the genes were co-transfected into HeLa cells with overexpression cassettes of either *SMARCAL1* or *BRG1*. As expected, transfection of the ShRNA construct against either the 3’ UTR of *SMARCAL1* or *BRG1* led to downregulation of *ATM* and *ATR* (Fig. 1M and N). This downregulation could be partially or completely reversed when cells were co-transfected with the overexpression construct of either *SMARCAL1* or *BRG1* respectively (Fig. 1M and N). Finally, ChIP confirmed the presence of both SMARCAL1 and BRG1 on *ATM* and *ATR* promoters (Supplementary Fig. 1C and D).

Thus, both *BRG1* and *SMARCAL1* appear to be co-regulating *ATM* and *ATR*. Downregulation of either *BRG1* or *SMARCAL1* results in transcription repression of *ATM* and *ATR*.

As both Sh*70* and Sh*71* cells behaved identically, we have used Sh*71* (termed as Sh*SMARCAL1*) in all the subsequent experiments.

### Downregulation of *SMARCAL1* and *BRG1* leads to impaired G2/M checkpoint trespass on DNA damage in HeLa cells

The phosphorylation of Chk2 and Chk1 by ATM/ATR kinases occurs only in the presence of DNA damage. To understand whether BRG1 and SMARCAL1 are important for activating the G2/M checkpoint on induction of DNA damage, the expression of BRG1, SMARCAL1, ATM and ATR was studied during DNA damage.

Previously, it has been shown that the expression of both SMARCAL1 and BRG1 is upregulated in HeLa cells on treatment with 2 μM doxorubicin for 10 minutes [36]. This upregulation of both SMARCAL1 and BRG1 was specific to doxorubicin-induced DNA damage and was not been observed with other DNA damaging agents [36,37].

Based on these data, we hypothesized that if SMARCAL1 and BRG1 are co-regulating *ATM* and *ATR*, then treatment with doxorubicin that results in upregulation of these two ATP-dependent chromatin remodeling proteins should also result in upregulation of these two DNA damage response genes.

As hypothesized, the expression of both *ATM* and *ATR* were indeed found to be upregulated in HeLa cells on treatment with 2 μM doxorubicin for 10 minutes (Fig. 2A-C).

**Figure 2.**
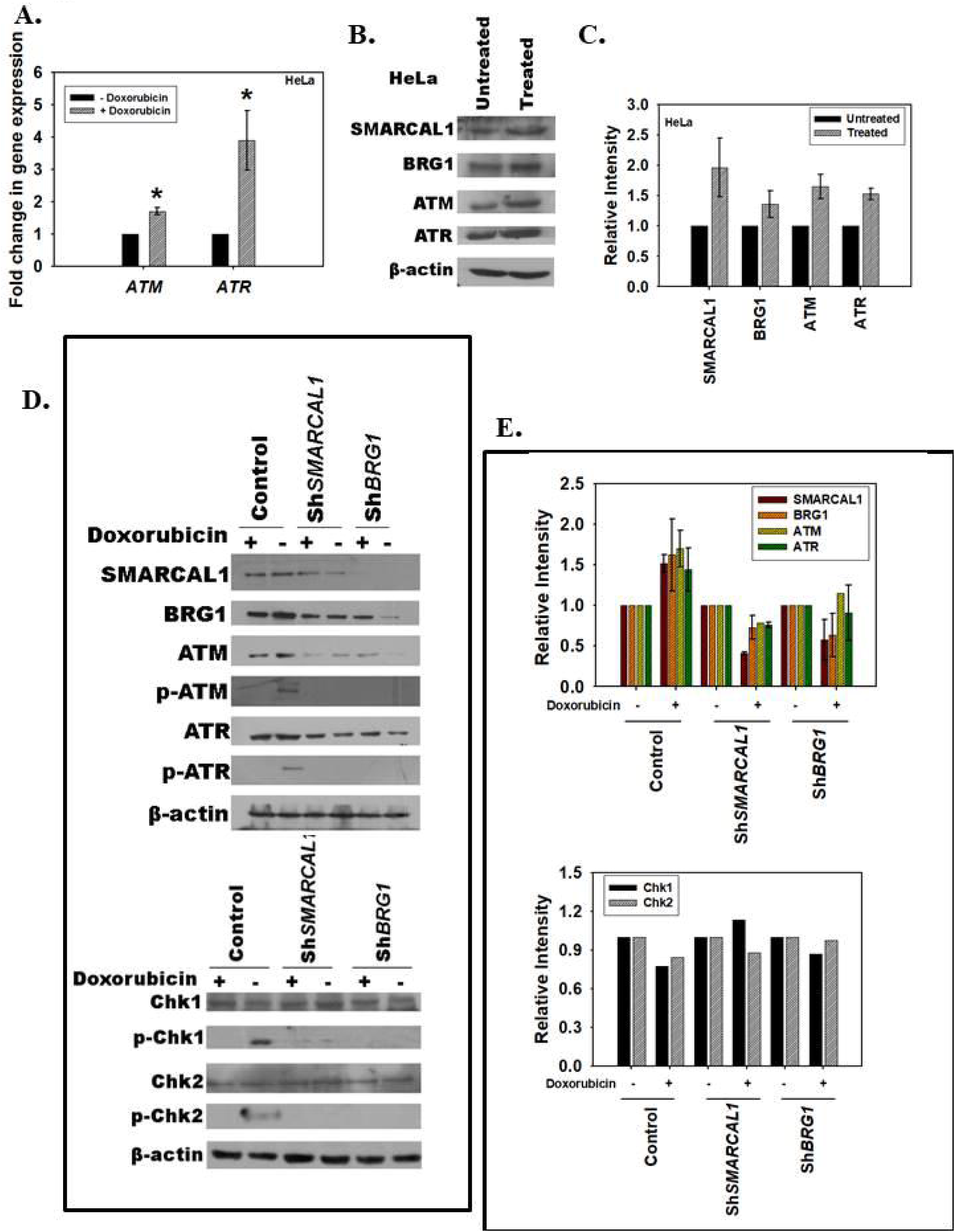
SMARCAL1 and BRG1 regulate the expression of ATM and ATR during doxorubicin-induced DNA damage. (A). The expression of *SMARCAL1*, *BRG1*, *ATM* and *ATR* was analysed in doxorubicin treated HeLa cells using qPCR. (B). The expression of SMARCAL1, BRG1, ATM and ATR was analysed in doxorubicin treated HeLa cells using western blot. (C). Quantitation of the western blot. (D). Western blot analysis of SMARCAL1, BRG1, ATM, p-ATM (Ser 1981), ATR, p-ATR (Ser 428), Chk1, p-Chk1 (Ser 345), Chk2 and p-Chk2 (Thr 68) in control (vector transfected), Sh*SMARCAL1*, and Sh*BRG1* cells in the absence and presence of doxorubicin. (E). Quantitation of the western blots. In these experiments, cells were treated with 2 μM doxorubicin for 10 minutes to induce DNA damage. The data in case of qPCR experiments is presented as average ±s.d of three independent experiments (*P < 0.001; unpaired student’s t-test).

Next the expression of SMARCAL1 and BRG1 along with the mediators of the G2/M checkpoint was studied in Sh*SMARCAL1* and Sh*BRG1* cells. It was found that in control cells (vector transfected), treatment with doxorubicin resulted in upregulation of SMARCAL1, BRG1, ATM, and ATR; however, in Sh*SMARCAL1* and Sh*BRG1* cells, the expression of these proteins were downregulated and on induction of DNA damage remained unchanged (Fig. 2D and E). Further, phosphorylation of ATM and ATR was also not detected in Sh*SMARCAL1* and Sh*BRG1* cells as compared to the control cells on doxorubicin treatment (Fig. 2D and E). Concomitantly, though the levels of Chk1 and Chk2 were unaltered in control, Sh*SMARCAL1* and Sh*BRG1* cells, the phosphorylation of Chk1 and Chk2 was impaired in the *SMARCAL1*/*BRG1* downregulated cells as compared to the control cells (Fig. 2D and E).

The loss of ATM and ATR, and therefore, impaired phosphorylation of Chk1 and Chk2, on doxorubicin treatment led to checkpoint trespass in Sh*SMARCAL1* and Sh*BRG1* cells as compared to the control cells as shown by β-tubulin staining (for mitotic spindles) and FACS. The control cells on doxorubicin treatment (as evidenced by the appearance of γH2AX foci) arrested at the G2/M checkpoint and none of the cells progressed into mitosis (as observed by the localization of β-tubulin to the cytoplasm and lack of mitotic spindles) (Fig. 3A-D). In contrast, 15% of the Sh*SMARCAL1* and Sh*BRG1* cells formed mitotic spindles indicating the progression into M phase (Fig. 3B and C). Further, FACS analysis showed a 3-fold increase in Sh*SMARCAL1/*Sh*BRG1* cells present in G2/M phase and a 2-fold increase in these cells present in G1 phase compared to the control cells (Fig. 3D).

**Figure 3.**
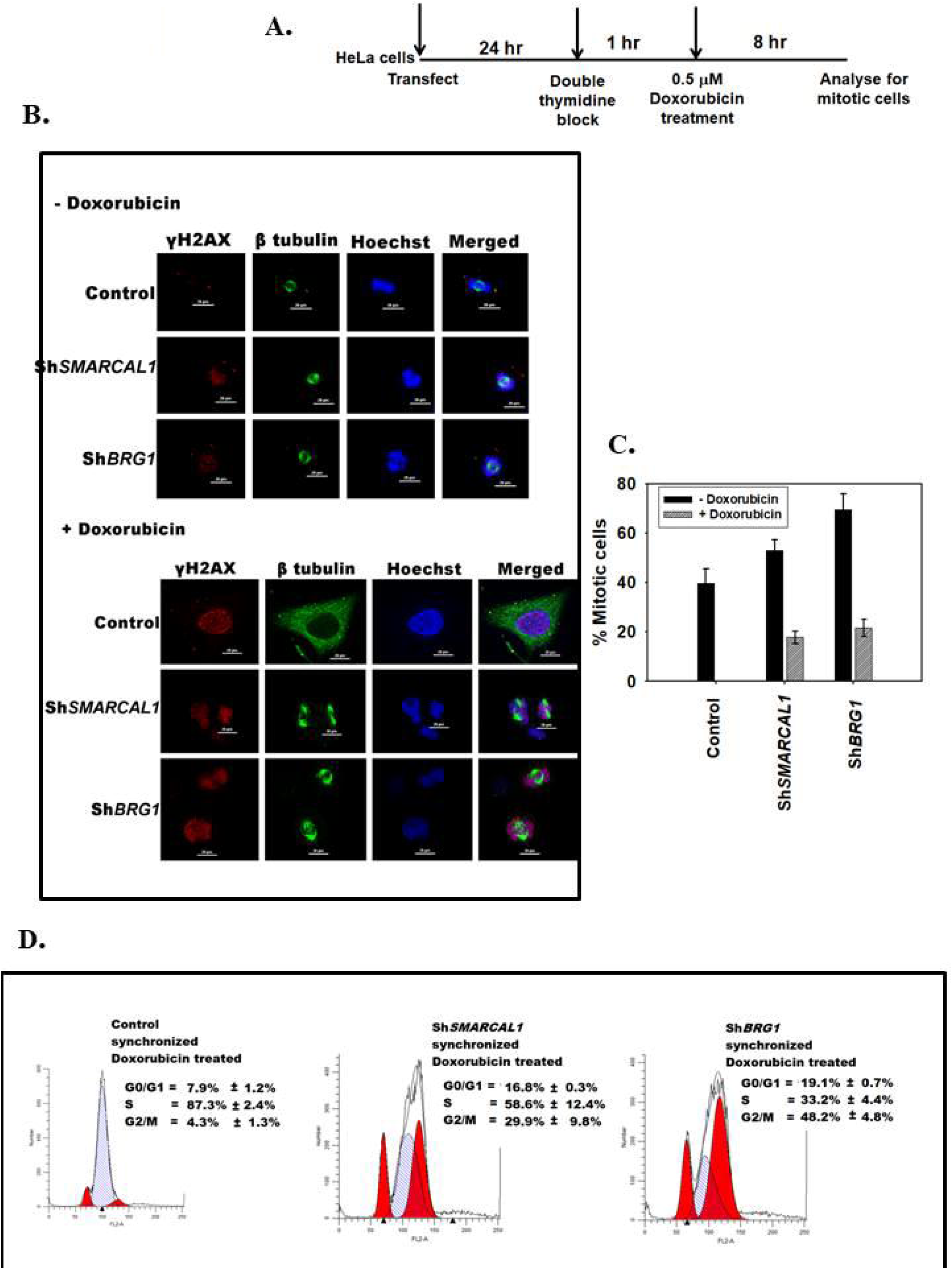
Downregulation of *SMARCAL1* and *BRG1* leads to impaired G2/M checkpoint trespass on DNA damage in HeLa cells. (A). Schematic representation showing the experimental methodology. Briefly, HeLa cells were transfected with vector alone, or with ShRNA against *SMARCAL1* or ShRNA against *BRG1*. The double thymidine block was started 24 hr after transfection. Cells were released from double thymidine block and 1 hr after release, were treated with 0.5 μM doxorubicin. Cells were harvested 8 hr later and analysed for the presence of mitotic cells either by immunostaining or by FACS. (B). The formation of γH2AX foci as well as spindle fibres (β-tubulin) in the absence and presence of doxorubicin was monitored in control (vector transfected), Sh*SMARCAL1* and Sh*BRG1* cells using immunostaining. (B). The number of mitotic cells (cells possessing spindle fibres) in the absence and presence of doxorubicin was quantitated in control (vector transfected), Sh*SMARCAL1* and Sh*BRG1* cells. The data is presented as an average ±s.d of two independent experiments (n ≥ 100 cells). (C). Analysis of cells in G1, S, G2/M stage of cell cycle was performed using FACS and the data is presented as average ±s.d of three independent experiments.

Thus, SMARCAL1 regulates the expression of *BRG1* and BRG1, in turn, regulates the expression of *SMARCAL1* creating a positive feedback loop in HeLa cells. Further, SMARCAL1 and BRG1 together positively co-regulate the expression of ATM and ATR in this cell line (Supplementary Fig. 2A).

### Regulation of *ATM/ATR* by SMARCAL1/BRG1 exists in other cell lines also

To determine whether this regulation operates in other cell lines expressing both SMARCAL1 and BRG1, the expression pattern of these two genes along with that of ATM and ATR was studied in HEK293 and MCF-7 cell lines. Downregulation of *SMARCAL1* in both HEK293 and MCF-7 led to downregulation of *BRG1* (Supplementary Fig. 2B and C). However, downregulation of *BRG1* in both these cell lines led to upregulation of *SMARCAL1*, indicating SMARCAL1 positively regulates the expression of BRG1 while BRG1 negatively regulates the expression of SMARCAL1 (Supplementary Fig. 2B and C). Further, downregulation of either *SMARCAL1* or *BRG1* led to upregulation of *ATM* and *ATR* in HEK293 cells; in MCF-7 cells, in contrast, both *ATM* and *ATR* were downregulated on downregulation of either *SMARCAL1* and *BRG1* (Supplementary Fig 2B and C).

Thus, in HEK293 cells, SMARCAL1 positively regulates BRG1 while BRG1 negatively regulates SMARCAL1; both negatively regulate ATM/ATR (Supplementary Fig 2D). In MCF-7cells, on the other hand, SMARCAL1 positively regulates BRG1 and BRG1 negatively regulates SMARCAL1; both positively regulate ATM/ATR (Supplementar Fig. 2E).

If this model is correct, then the upregulation of *SMARCAL1*, *BRG1*, *ATM* and *ATR* should not be observed with doxorubicin-induced DNA damage in MCF-7 cells. The control (vector transfected) MCF-7 cells were treated with 2 μM doxorubicin for 10 minutes and as hypothesized, the expression of the four genes did not alter with doxorubicin-induced DNA damage (Supplementary Fig. 2F).

Thus, a regulatory loop does exist between SMARCAL1, BRG1, ATM and ATR; however, the nature of the loop varies from cell line to cell line.

### Both SMARCAL1 and BRG1 are required for the upregulation of *ATM* and *ATR* on doxorubicin-induced DNA damage

From the above experiments, though it is evident that both SMARCAL1 and BRG1 regulate ATM/ATR, it is not clear whether both are necessary or whether one remodeler is sufficient. As it is technically not possible to create a HeLa cell where only *SMARCAL1* or *BRG1* is downregulated, cell lines that lacked either SMARCAL1 (HepG2 cell line) or BRG1 (A549 cell line) were used to understand whether both the proteins were required for the upregulation of ATM and ATR expression on doxorubicin treatment. However, it needs to be pointed, given the variances between the cell lines (as outlined in section 3.4), the differences observed in A549 and HepG2 do not offer a complete representation of the role of SMARCAL1 and BRG1 in regulating *ATM* and *ATR*.

In both A549 and HepG2 cell lines, *ATM* and *ATR* were downregulated on treatment with 2 μM doxorubicin for 10 minutes; however, the protein levels were not significantly altered (Supplementary Fig. 3A-F) indicating both SMARCAL1 and BRG1 are required for transcriptional regulation of these genes. However, in these cells, unlike the Sh*SMARCAL1*/Sh*BRG1* cells, ATM and ATR are phosphorylated on DNA damage indicating that the DNA damage response is not altered in these cell lines (Supplementary Fig. 3G). Thus, it appears that the phosphorylation of ATM and ATR is not dependent on their upregulation on DNA damage by SMARCAL1/BRG1; however, if ATM and ATR are repressed as in *SMARCAL1/BRG1*downregulated HeLa cells, phosphorylation is affected and hence, the phosphorylation of downstream effector molecules is impaired leading to non-functional G2/M checkpoint.

### SMARCAL1 and BRG1 recruit RNAPII to the promoters of *ATM* and *ATR*

These three cell lines-HeLa, A549, and HepG2- were now used delineate the mechanism of transcription regulation of *ATM* and *ATR* by analysing the occupancy of SMARCAL1, BRG1, RNAPII and H3K9Ac, an activation mark [46], on these promoters.

ChIP experiments showed that on doxorubicin treatment, in HeLa cells, the occupancy of BRG1, SMARCAL1, RNAPII, and H3K9Ac increased upstream (primer pair 2) of the transcription start site (TSS) of the *ATM* promoter (Fig. 4A). Downstream of the TSS (primer pair 3), the occupancy of BRG1 and RNAPII did not change while the occupancy of H3K9Ac increased. ChIP-reChIP experiment showed that on doxorubicin treatment BRG1 and SMARCAL1 were present simultaneously on the *ATM* promoter at the region amplified by primer pair 2 (Supplementary Fig. 4A). SMARCAL1 is known to bind to DNA regions possessing secondary structures. MFold [47] and QGRS [48] showed that the region amplified by primer pair 2 can indeed form secondary structures (Supplementary Table 1). Using ADAAD, the bovine homolog of SMARCAL1, it was confirmed that the protein was indeed able to bind to the *ATM* promoter primer pair 2 region *in vitro* (Supplementary Fig. 4B). Previously, using CD spectroscopy [49], ADAAD has been shown to remodel promoter DNA [36–38]. Using the same technique we confirmed that ADAAD can indeed bind to the ATM promoter and remodel DNA (Supplementary Fig. 4C).

**Figure 4.**
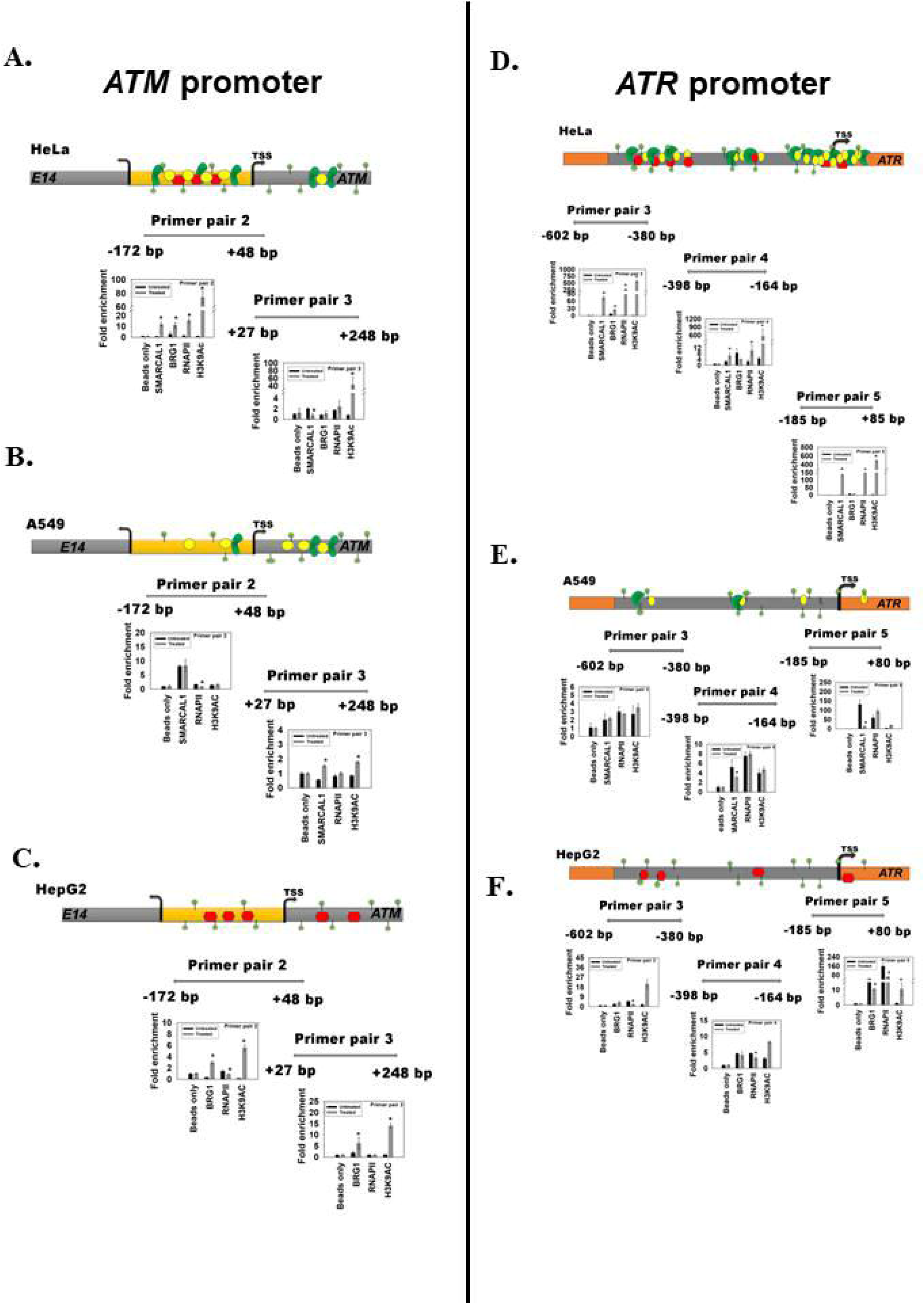
The occupancy of SMARCAL1 and BRG1 is altered on the *ATM* and *ATR* promoter on doxorubicin treatment. The occupancy of SMARCAL1, BRG1, RNAPII and H3K9Ac was monitored on the *ATM* promoter on doxorubicin treatment in (A). HeLa cells; (B). A549 cells; and in (C) HepG2 cells. The occupancy of SMARCAL1, BRG1, RNAPII and H3K9Ac was monitored on the *ATR* promoter on doxorubicin treatment in (D). HeLa cells; (E). A549 cells; and in (F) HepG2 cells. The data all these experiments is presented as average ±s.d of three independent experiments (*P < 0.001; unpaired student’s t-test).

In A549 cells, where BRG1 is not expressed, the occupancy of SMARCAL1 and H3K9Ac did not change on doxorubicin treatment upstream of the TSS while the occupancy of RNAPII decreased correlating with decreased transcript levels indicating BRG1 is needed for recruiting these three proteins to the promoter region (Fig. 4B). In HepG2 cells, where SMARCAL1 is not expressed, the RNAPII occupancy decreased on doxorubicin treatment upstream of the TSS correlating with decreased expression (Fig. 4C). However, BRG1 and H3K9Ac occupancy increased indicating SMARCAL1 is needed possibly only for the recruitment of RNAPII (Fig. 4C).

On *ATR* promoter, ChIP experiments showed that in HeLa cells the occupancy of BRG1, SMARCAL1, RNAPII, and H3K9Ac increased both upstream and downstream of the TSS on doxorubicin treatment (Fig. 4D). ChIP-reChIP experiment confirmed that BRG1 and SMARCAL1 are present simultaneously on the *ATR* promoter (primer pair 3) on doxorubicin treatment (Supplementary Fig. 4D). Analysis using MFold and QGRS showed that the promoter region amplified by primer pair 5 has the potential to form secondary structures (Supplementary Table 1). Using ADAAD, the bovine homolog of SMARCAL1, it was confirmed that the protein can bind to the *ATR* promoter primer pair 5 region *in vitro* (Supplementary Fig. 4E). Further, using CD spectroscopy, it was confirmed that the protein can remodel the promoter DNA (Supplementary Fig. 4F).

In A549 cells, the occupancy of SMARCAL1, RNAPII, and H3K9Ac was unaltered upstream of the TSS (primer pair 3 and 4) while the occupancy of SMARCAL1 decreased around the TSS (primer pair 5), indicating BRG1 is needed for recruiting SMARCAL1, RNAPII, and H3K9Ac to the *ATR* promoter on doxorubicin treatment (Fig. 4E). In HepG2 cells, the occupancy of BRG1 on the *ATR* promoter was unaltered on doxorubicin treatment but the occupancy of H3K9Ac increased upstream as well as around the TSS (Fig. 4F). The occupancy of RNAPII decreased both upstream as well as around the TSS (Fig. 4F), indicating once again that SMARCAL1 is needed for RNAPII recruitment.

Thus, based on these results we propose that both BRG1 and SMARCAL1 are needed for recruitment of RNAPII to the *ATM* and *ATR* promoters on doxorubicin-induced DNA damage; in addition, BRG1 possibly helps in recruiting SMARCAL1 to these promoters.

### Phosphorylation of ATM and ATR is necessary for the activation of the transcriptional network

The experimental results presented till now show that SMARCAL1 and BRG1 co-regulate the expression of *ATM* and *ATR*. Previously, SMARCAL1 and BRG1 had been shown to co-regulate the RNAi genes-*DROSHA*, *DGCR8*, and *DICER-* on doxorubicin treatment, and thereby regulating the formation of ncRNA required for 53BP1 foci formation [37]. Therefore, we propose that in HeLa cells a transcriptional network exists wherein SMARCAL1 and BRG1 transcriptionally regulate DNA damage response genes-*ATM*, *ATR*, *DROSHA*, *DGCR8*, and *DICER*- and hence, regulate the response of the cell to DNA damage agents.

Reports have also shown that the activity of both SMARCAL1 and BRG1 are regulated by phosphorylated ATR and ATM respectively [22,34]. Therefore, the importance of phosphorylated ATM and ATR in this transcriptional network was next investigated. HeLa cells were treated with either ATM inhibitor (ATMi) or ATR inhibitor (ATRi) for 24 hours prior to treatment with 2 μM doxorubicin for 10 minutes. Treatment with ATMi inhibited ATM phosphorylation while ATRi treatment inhibited Chk1 phosphorylation, confirming that the inhibitor treatment functioned as reported previously [11,17,50–52] (Fig. 5A). Quantitative PCR showed that neither *SMARCAL1* nor *BRG1* were upregulated when cells were treated with ATMi/ATRi and doxorubicin (Fig. 5B and C). As a consequence, upregulation of *ATM, ATR, DROSHA, DGCR8,* and *DICER* was also not observed in these cells (Fig. 5B and C). This was confirmed by western blots (Fig. 5A; quantitation of blots is shown in Supplementary Fig. 5A).

**Figure 5.**
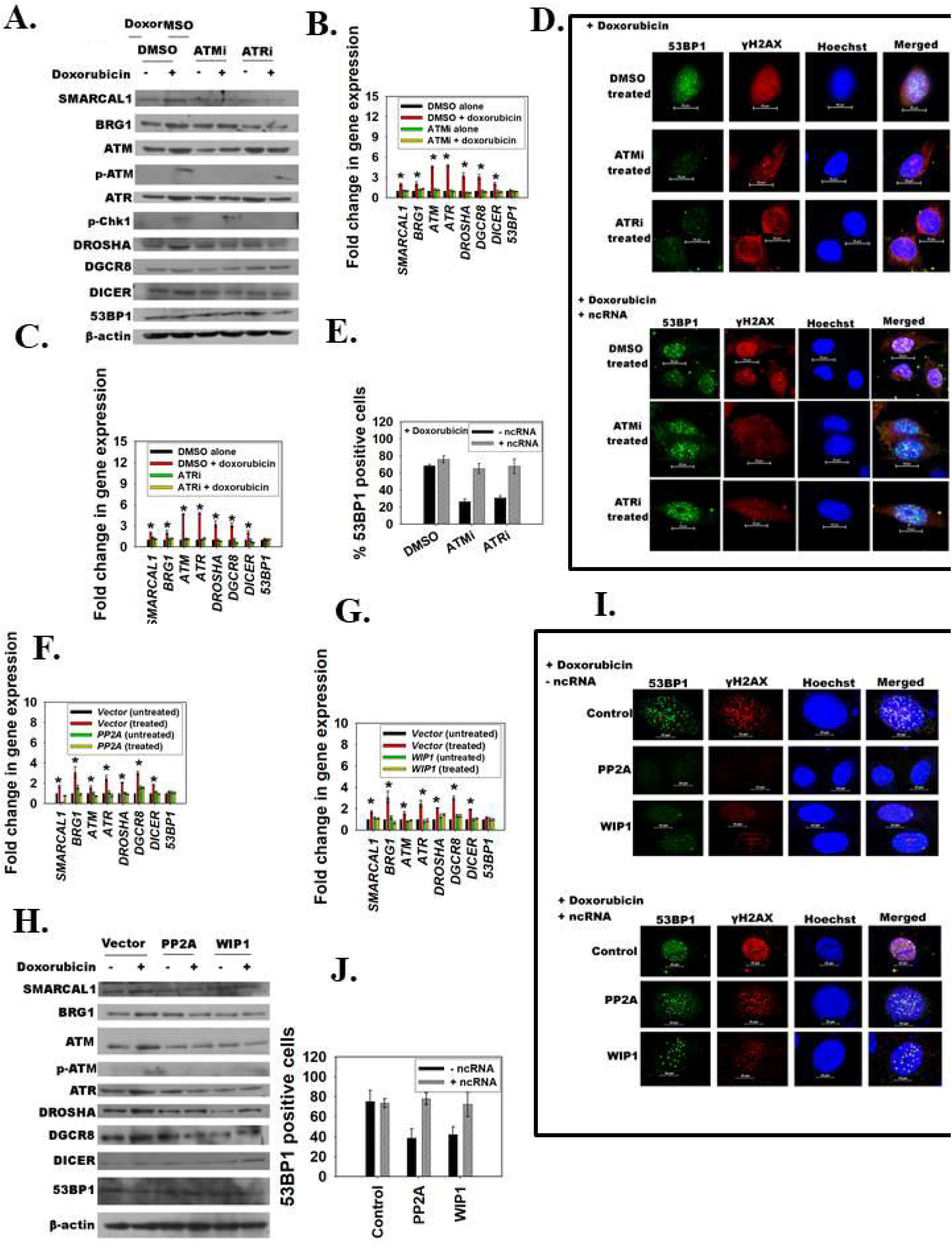
Phosphorylation of ATM and ATR feedback regulates transcription of*ATM* and *ATR* by SMARCAL1 and BRG1. (A). Protein expression after treatment with doxorubicin was analysed in absence and presence of ATMi and ATRi using western blot. (B). Expression of the genes in the presence of ATMi. (C). Expression of the genes in the presence of ATRi. (D). The formation of 53BP1 and γH2AX foci in the presence of ATMi and ATRi was analysed using immunofluorescence before and after addition of ncRNA purified from doxorubicin treated HeLa cells. (E). The number of cells (n≥100) containing 53BP1 foci (≥5foci per cell) were quantitated in cells (untreated, ATMi treated, and ATRi treated) before and after adding ncRNA. (F). Expression of genes in HeLa cells transfected with *PP2A* gene in the absence and presence of doxorubicin. (G). Expression of genes in HeLa cells transfected with *WIP1* gene in the absence and presence of doxorubicin. (H). Analysis of protein expression using western blots. (I). The formation of 53BP1 foci was analysed in HeLa cells transfected either with vector alone, or *PP2A* gene or *WIP1* gene in the absence and presence of ncRNA after treatment with 2 μM doxorubicin for 10 min. (J). The number of cells (n≥ 100) containing 53BP1 foci (≥ foci per cell) was quantitated in cells (transfected either with vector alone, *PP2A*, or *WIP1)*. ncRNA used in these experiments was purified from HeLa cells treated with 2 μM doxorubicin for 10 min. The data in case of qPCR experiments is presented as average ±s.d of three independent experiments (*P < 0.001; unpaired student’s t-test).

As stated earlier, the upregulation of the three RNAi genes by SMARCAL1 and BRG1 is needed for 53BP1 foci formation [37]. Therefore, in cells treated with ATM and ATR inhibitors, 53BP1 foci formation should be abrogated on doxorubicin treatment. Experimental results showed that indeed in cells treated with ATM and ATR inhibitors, 3-fold less cells formed 53BP1 foci as compared to control cells on doxorubicin treatment (Fig. 5D and E; Control experiments are shown in Supplementary Fig. 5B). The formation of 53BP1 foci was restored on addition of ncRNA from HeLa cells treated with doxorubicin (Fig. 5D and E). It needs to be pointed out that the expression of 53BP1 was not changed when cells were treated with ATM and ATR inhibitor (Fig. 5A-C; quantitation of blots is shown in Supplementary Fig. 5A).

Two phosphatases- PP2A and WIP1-have been identified to play a key role in dephosphorylating ATM [13,14]. If phosphorylation of ATM is important for the transcriptional network mediated by SMARCAL1 and BRG1, then overexpression of WIP1 and PP2A should result in abrogation of the network and therefore, of 53BP1 foci formed at the DNA damage site.

*PP2A-A* (the regulatory subunit; henceforth, referred as *PP2A*) and *WIP1* genes were transfected into HeLa cells prior to treatment with 2 μM doxorubicin for 10 minutes. The expression of SMARCAL1 and BRG1 were found to be unchanged on doxorubicin treatment in cells transfected with either *PP2A* or *WIP1* (Fig. 5F and G). This was corroborated by the western blots (Fig. 5H; quantitation of blots is shown in Supplementary Fig. 5C). As expected, the downstream genes-*ATM*, *ATR*, *DROSHA*, *DGCR8*, and *DICER*- were also unchanged on doxorubicin treatment in cells transfected with either *PP2A* or *WIP1* (Fig. 5F-H; quantitation of blots is shown in Supplementary Fig. 5C). Finally, 53BP1 foci formation was also found to be abrogated in cells transfected with either *PP2A* or *WIP1* (Fig. 5I and J; Control experiments are shown in Supplementary Fig. 5D). The 53BP1 foci formation, once again, could be rescued by addition of ncRNA from HeLa cells treated with doxorubicin (Fig. 5I and J).

### Dephosphorylation of ATM and ATR results in cessation of the DNA damage response

Based on the above results, it was hypothesized that the feedback regulation of the transcriptional loop by ATM and ATR is needed to amplify the DNA damage response signal and dephosphorylation of ATM and ATR would lead to cessation of this signal, thus, possibly bringing the DNA damage response pathway to a halt. To test the hypothesis, HeLa cells were treated with okadiac acid, the inhibitor of PP2A along with doxorubicin. Cells were released from doxorubicin treatment and the expression of SMARCAL1, BRG1, ATM and ATR was monitored. It was observed that the expression levels of SMARCAL1, BRG1, ATM and ATR were upregulated on doxorubicin treatment for 10 minutes (corresponding to 0 min time after release) both in the absence and presence of okadiac acid (Fig. 6A-H). In the absence of doxorubicin, the expression of SMARCAL1, BRG1, ATM and ATR returned back to normal levels within 5 min of removal of doxorubicin (Fig. 6A-E; G-J). Further, ATM was also dephosphorylated within 5 minutes (Fig. 6E). However, in the presence of okadiac acid, the expression of these genes remain upregulated and return to normal only around 30 min after removal of doxorubicin (Fig. 6A-D; F-J) corresponding with ATM remaining in the phosphorylated form (Fig. 6F).

**Figure 6.**
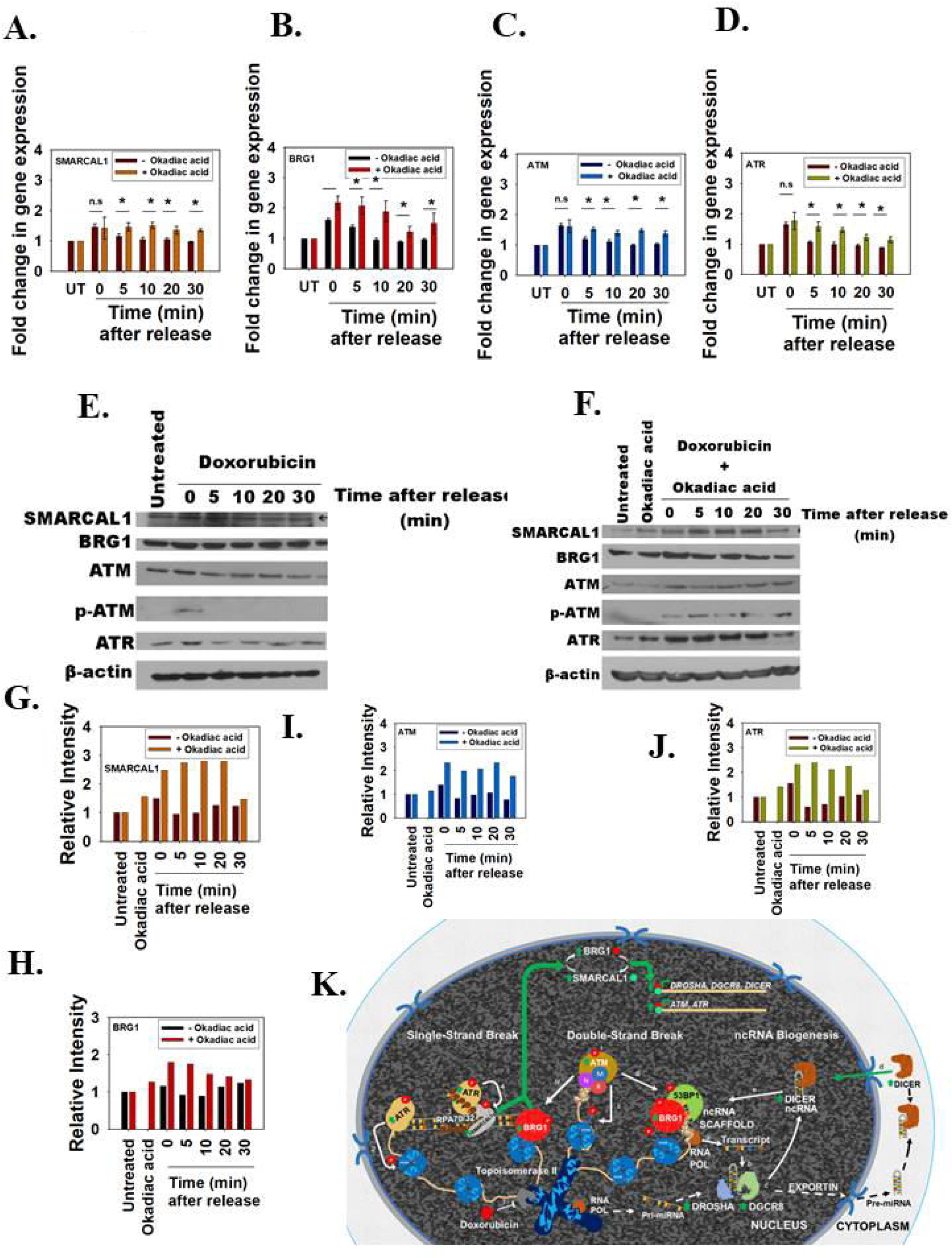
Dephosphorylation of ATM and ATR results in cessation of the DNA damage response. HeLa cells were treated with okadiac acid followed by 2 μM doxorubicin for 10 min after which doxorubicin-containing media was removed and fresh media was added. Cells were harvested at the time points indicated and expression was analysed by qPCR for (A). *SMARCAL1*; (B). *BRG1*; (C). *ATM*; (D). *ATR*. (E). Protein expression was analysed using western blot in cells after release from doxorubicin treatment. (F). Protein expression was analysed using western blot in okadiac acid treated cells after release from doxorubicin treatment. Quantitation of the western blots for (G). SMARCAL1; (H). BRG1; (I). ATM; (J). ATR. (K). Model explaining the transcriptional regulation of G2/M checkpoint. *SMARCAL1* and *BRG1* are mutually co-regulated in HeLa cells. Together they co-regulate the expression of *ATM*, *ATR*, *DROSHA*, *DGCR8*, and *DICER*. On induction of DNA damage by doxorubicin, the expression of all these genes is upregulated and this modulation is mediated by SMARCAL1 and BRG1. The signal remains in ON state due to phospho-ATM and phospho-ATR. The result is the formation of ncRNA that enables the formation of 53BP1 foci at the site of DNA damage. Dephosphorylation of ATM and ATR leads to cessation of the signal. The data in case of qPCR experiments is presented as average ±s.d of three independent experiments (*P < 0.001; unpaired student’s t-test).

Thus, based on these results, we propose that on DNA damage in HeLa cells, SMARCAL1 and BRG1 are upregulated. This upregulation mediates transcriptional activation of DNA damage response genes-*ATM* and *ATR-* as well as the three RNAi genes-*DROSHA*, *DGCR8*, and *DICER*-leading to the formation of 53BP1 foci. This transcriptional loop is kept in active (switched ON state) mode by phospho-ATM and phospho-ATR. Dephosphorylation of these kinases results in cessation of the signal (switched OFF state) (Fig. 6K).

### The mutations present in SIOD and CSS patients lead to abrogation of DNA damage response signal

Mutations in SMARCAL1 are associated with Schimke Immunoosseous Dysplasia (SIOD) while mutations in BRG1 are associated with Coffin-Siris Syndrome (CSS) [27,35]. We rationalized that if SMARCAL1 and BRG1 are necessary for DNA damage response pathway, then the patient-associated mutations should abrogate the DNA damage response. Therefore, the role of three SIOD-associated mutants-A468P, I548N, and S579L- was studied as previously it has been shown that these three mutants are unable to hydrolyze ATP due to impaired DNA binding [53]. In addition, three CSS-associated mutants-T895M, L921F, and M1011T-were also chosen as these three mutants mapped to the ATPase domain and are postulated to lack ATPase activity [27].

HeLa cells were co-transfected with Sh*RNA* construct targeting the 3′ UTR of *SMARCAL1* along with a plasmid either containing the gene for full-length *SMARCAL1* or containing the gene encoding a SIOD-associated mutation or containing the gene encoding for an ATPase dead mutant of SMARCAL1 called K464A [53]. Co-transfection of Sh*RNA*(*SMARCAL1*) along with full-length *SMARCAL1* resulted in upregulation of *SMARCAL1*, *BRG1*, *ATM* and *ATR* genes on doxorubicin treatment (Fig. 7A). This upregulation was not observed when Sh*RNA(SMARCAL1)* was co-transfected along with K464A mutant or SIOD-associated mutants indicating that the ATPase activity of SMARCAL1 is important for the transcriptional upregulation (Fig. 7A).

**Figure 7.**
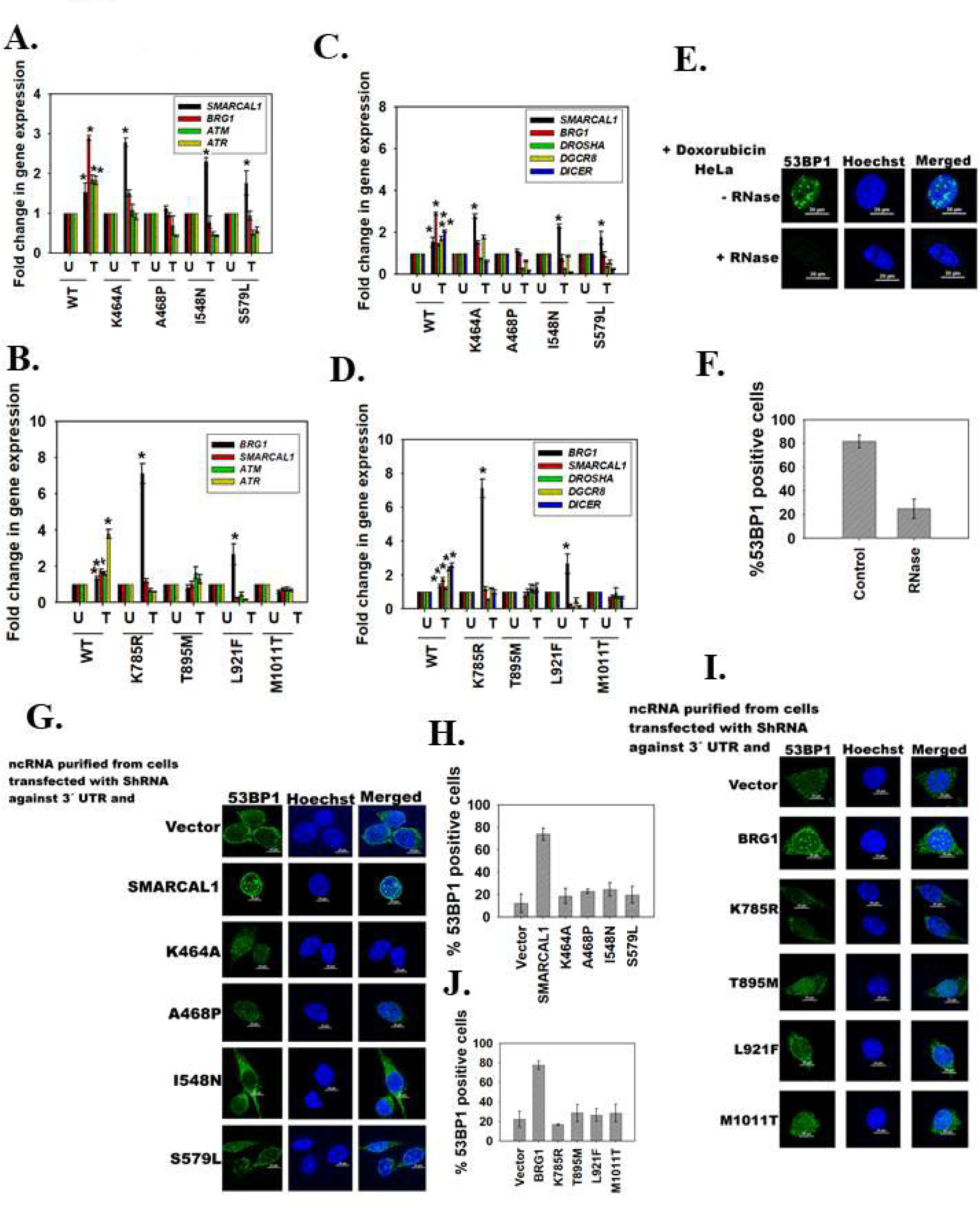
Mutations present in SIOD and CSS patients lead to abrogation of DNA damage response signal. (A). Expression of *SMARCAL1*, *BRG1*, *ATM* and *ATR* was analysed after co-transfection of HeLa cells with Sh*RNA*(*SMARCAL1)* along with vector alone or vector containing wild type *SMARCAL1*, or *K464A* mutant, or *A468P* mutant, or *I548N*, or *S579L* mutant gene. The data is presented as average ±s.d of two independent experiments (*P < 0.001; unpaired student’s t-test). (B). Expression of *SMARCAL1*, *BRG1*, *ATM* and *ATR* was analysed after co-transfection of HeLa cells with Sh*RNA*(*BRG1)* along with vector alone or vector containing wild type *BRG1*, or *K785R* mutant, or *T895M* mutant, or *L921F*, or *M1011T* mutant gene. The data is presented as average ±s.d of two independent experiments (*P < 0.01; unpaired student’s t-test). (C). Expression of *SMARCAL1*, *BRG1*, *DROSHA, DGCR8* and *DICER* was analysed after co-transfection of HeLa cells with Sh *RNA(SMARCAL1)* along with vector alone or vector containing wild type *SMARCAL1*, or *K464A* mutant, or *A468P* mutant, or *I548N*, or *S579L* mutant gene. The data is presented as average ±s.d of two independent experiments (*P < 0.001; unpaired student’s t-test). (D). Expression of *SMARCAL1*, *BRG1*, *DROSHA*, *DGCR8* and *DICER* was analysed after co-transfection of HeLa cells with Sh*BRG1* along with vector alone or vector containing wild type *BRG1*, or *K785R* mutant, or *T895M* mutant, or *L921F*, or *M1011T* mutant gene. The data is presented as average ±s.d of two independent experiments (*P < 0.05; unpaired student’s t-test). (E). The formation of 53BP1 foci in HeLa in the absence and presence of RNase was monitored after treatment with 2 μM doxorubicin for 10 min. (F). The number of cells (n≥ 0; 2 independent experiments) containing 53BP1 foci (≥5) was quantitated in HeLa cells in the absence and presence of RNase after treatment with 2 μM doxorubicin for 10 min. (G). The restoration of 53BP1 foci in HeLa cells on doxorubicin and RNase treatment was monitored in the presence of ncRNA purified from co-transfected (Sh*SMARCAL1* along with vector containing genes either for wild type *SMARCAL1*, or K464A, or A468P, or I548N, or S579L gene) HeLa cells. (H). The number of cells (n≥100; 2 independent experiments) containing 53BP1 foci (≥5) was quantitated for the images shown in Fig. 7G. (I). The restoration of 53BP1 foci in HeLa cells on doxorubicin and RNase treatment was monitored in the presence of ncRNA purified from co-transfected (Sh*BRG1* along with vector containing genes either for wild type *BRG1*, or *K785R*, or *T895M,* or L921F, or M1011T gene) HeLa cells. (J). The number of cells (n≥100; 2 independent experiments) containing 53BP1 foci (≥5) was quantitated for the images shown in Fig. 7I.

A similar observation was made in case of CSS-associated mutants. When HeLa cells were co-transfected with Sh*RNA* construct targeting the 3’ UTR of *BRG1* along with a plasmid encoding for full-length wild type *BRG1*, the expression of *SMARCAL1*, *BRG1*, *ATM* and *ATR* were upregulated on doxorubicin treatment (Fig.7B). However, when S*hRNA(BRG1)* construct was co-transfected with ATPase dead mutant of BRG1, K785R, or with genes encoding for CSS-associated mutants, the expression of *SMARCAL1*, *BRG1*, *ATM* and *ATR* were not upregulated on doxorubicin treatment (Fig. 7B).

Next, the ability of SIOD-associated and CSS-associated mutants to form non-coding RNA required for the formation of 53BP1 foci formation was assessed. Co-transfection of Sh*RNA* (*SMARCAL1)* construct with wild type *SMARCAL1* gene resulted in upregulation of *DROSHA*, *DGCR8*, and *DICER* expression on doxorubicin treatment (Fig. 7C). A similar result was observed when HeLa cells were co-transfected with Sh*RNA*(*BRG1)* construct along with either wild type *BRG1* gene or with K785R mutant or with CSS-associated mutants (Fig. 7D). To estimate the ability of the mutants to support the formation of 53BP1 foci, non-coding RNA was purified from these co-transfected cells. HeLa cells were treated with 2 μM doxorubicin for 10 minutes leading to the formation of 53BP1 foci (Fig. 7E and F). RNase treatment, as expected, resulted in abrogation of these foci (Fig. 7E and F). RNase was inactivated by treatment with RNase inhibitor and the cells were incubated with ncRNA purified from the co-transfected cells. Non-coding RNA purified from cells co-transfected with Sh*RNA* and wild-type protein was able to restore the formation of 53BP1 foci (Fig 7G-J). However, ncRNA purified from cells co-transfected with ShRNA along with mutant protein was unable to restore formation of 53BP1 foci (Fig. 7G-J) indicating DNA damage response is affected in cells containing SIOD-associated SMARCAL1 mutations or CSS-associated BRG1 mutations.

## DISCUSSION

The G2/M cell cycle checkpoint mediated by ATM and ATR kinases is important for ensuring that the cell gets ample time to effect repair of the damaged DNA, and thus, helps in maintaining the fidelity of chromosome segregation during cell division [42,54]. The proteins involved in this checkpoint and the role of post-transcriptional modifications in effecting the repair have been well-characterized. However, very few reports exist for transcriptional regulation of *ATM* and *ATR*. Inhibition of Nuclear factor like-2 (NRF2), a transcription factor, has been reported to cause transcriptional repression of *ATM* and *ATR* [55]. Peng et al. have reported that downregulation of DNA-PKcs causes transcriptional repression of *ATM* [56]. Histone deacetylases have also been reported to transcriptionally regulate *ATM* [57]. However, none of these reports have characterized the mechanism regulating the transcription of these genes.

In this paper, the existence of a transcriptional regulatory loop controlling the expression of *ATM* and *ATR* and thus, the G2/M checkpoint in mammalian cells has been delineated. This transcriptional regulatory loop is mediated by the action of two transcriptional co-regulators, BRG1 and SMARCAL1, belonging to the family of ATP-dependent chromatin remodeling proteins [58].

BRG1 and SMARCAL1 mutually co-regulate each other [36]; thus, downregulation of *BRG1* results in reduced *SMARCAL1* levels and downregulation of *SMARCAL1* leads to repression of *BRG1* expression in HeLa cells. Further, downregulation of *BRG1*/*SMARCAL1* results in mitotic abnormalities and an ability to override the G2/M checkpoint as these two proteins drive the transcription of *ATM* and *ATR*. As in the case of the RNAi genes (*DROSHA*, *DGCR8* and *DICER*) [37], BRG1 and SMARCAL1 together co-regulate the transcription of these two kinases.

The transcriptional activation of *ATM* and *ATR* by SMARCAL1 and BRG1 is important for mediation of DNA damage response. On doxorubicin-induced DNA damage, both BRG1 and SMARCAL1 are upregulated in HeLa cells. This upregulation, in turn, mediates transcriptional activation of ATM and ATR as well as of the three RNAi genes. This leads to the activation of the G2/M checkpoint as well as synthesis of ncRNA required for 53BP1 foci formation as a part of DNA damage response. To keep this transcriptional control in an active mode (switch ON), phosphorylation of ATM and ATR is needed. Dephosphorylation of these two kinases results in shutting down of the transcriptional loop (switch OFF) (Fig. 6K). Both BRG1 and SMARCAL1 are known to be phosphorylated by ATM and ATR respectively [22,34]. Thus, it is possible that phosphorylated BRG1 and SMARCAL1 act as a transcriptional co-regulator on DNA damage. However, the mechanism of feedback regulation needs to be further delineated.

The ATPase activity of both BRG1 and SMARCAL1 is needed for the activation of the transcriptional loop. Thus, SIOD-associated mutants and CSS-associated mutants that cannot hydrolyse ATP are unable to mount an appropriate DNA damage response, indicating a possible DNA damage response defect in these patients.

Mechanistically, both SMARCAL1 and BRG1 are required for recruiting RNAPII to the *ATM* and *ATR* promoter. Absence of either protein results in downregulation of *ATM* and *ATR* on doxorubicin treatment, as in the case of A549 and HepG2 cells. Further, transcriptional regulation of ATM and ATR, and therefore, the G2/M checkpoint, appears to exist in all cells that contain both SMARCAL1 and BRG1, though the nature of the transcriptional regulatory loop varies from cell line to cell line. Thus, in HeLa cells, SMARCAL1 and BRG1 regulate each other and co-regulate *ATM* and *ATR* in a positive manner while in MCF-7 and HEK293 cells, SMARCAL1 positively regulates *BRG1* but BRG1 negatively regulates *SMARCAL1.* SMARCAL1 and BRG1 positively co-regulate *ATM* and *ATR* in MCF-7 cells while in HEK293, these two proteins negatively co-regulate *ATM* and *ATR*. As transcriptional regulation requires the action of transcription factors in addition to co-regulators such SMARCAL1 and BRG1, it is possible that the variances observed between different cell lines are due to different transcription factors. This needs to be further elucidated.

ATM and ATR kinases regulate not only the G2/M checkpoint but also control centrosome biogenesis as well as the spindle assembly checkpoint. Both the kinases have been shown to localize to the centrosome during mitosis [59]. Downregulation of *ATR* using siRNA results in centrosome amplification, and therefore, formation of multipolar spindles [60]. ATM controls the PARylation of the spindle pole organizer protein, NuMA1 by phosphorylating the protein at positions S1262 and S1601 [61]. As PARylation of NuMA1 is needed for correct spindle assembly, treatment of HeLa cells with ATM inhibitor, KU-55933, has been shown to result in multipolar spindles [61]. ATM is also required for localization of MDC1 to the kinetochores and thus, for activation of the spindle assembly checkpoint [62]. Thus, the mitotic defects seen on downregulation of SMARCAL1/BRG1 in HeLa cells could be due to transcriptional repression of *ATM* and *ATR* kinases.

The results presented in this paper augment the role of BRG1 and SMARCAL1 in maintaining genomic stability. Both these proteins mediate DNA repair and the genomic instability in absence of either of these proteins has been attributed to the accumulation of damaged DNA. Our results indicate that in addition to a direct role in DNA repair these two proteins are also responsible for mounting an appropriate DNA damage response and activating the G2/M cell cycle checkpoint via a transcriptional loop involving the ATM and ATR kinases as well as the RNAi genes.

## MATERIALS AND METHODS

### Reagents

Reagents for cell culture were purchased from HiMedia, USA. Sodium bicarbonate, TRI reagent, Hoechst 33342, doxorubicin, ATM inhibitor (KU-55933), ATR inhibitor (NU6027), thymidine, and propidium iodide was purchased from Sigma-Aldrich, USA. Molecular biology reagents were purchased either from MBI Fermentas (USA) or from NEB (USA). Protein-G fast flow bead resin was purchased from Merck-Millipore (USA). Micro-amp Fast 96-well reaction plates (0.1 ml) and micro-amp optical adhesive films were purchased from Applied Biosystems (USA). 2X fast SYBR Green PCR master mix was purchased from Kapa Biosystems (USA). For western blotting, Immobilon-P PVDF membrane was purchased from Merck-Millipore (USA).

### Antibodies

Antibodies to ATM (1:500 dilution; Catalog # D2E2), phospho-ATM (Ser 1981) (1:500 dilution; Catalog # 13050S), ATR (1:500 dilution; Catalog # E1S3S), phospho-ATR (Ser 428) (1: 500 dilution; Catalog #2853S) and RNAPII (1:1000 dilution; Catalog # 2629S) were purchased from Cell Signaling Technology (USA). Antibodies to BRG1 (1:1000 dilution; Catalog #B8184), H3K9Ac (Catalog #H0913), γH2AX (Catalog #H5912), β-actin (1:3000 dilution; Catalog #A1978), Chk1 (1:100; Catalog # SAB4500208), phospho-Chk2 (Thr 68) (1:500; Catalog # SAB4504367), Chk2 (1:1000; Catalog # C9233), CDK1 (1:1000; Catalog # HPA003387), and β-tubulin (Catalog # T8328) were purchased from Sigma-Aldrich (USA). Phospho-Chk1 (Ser 345) (1:200 dilution; Catalog # sc-17922) was purchased from Santa Cruz Biotechnology (USA). SMARCAL1 antibody was raised against N-terminus HARP domain by Merck (India) (Catalog # 106014) [36,41,46]. The TRITC- and FITC-conjugated anti-rabbit (1:100 dilution; Catalog # RTC2) and anti-mouse (1:100 dilution; Catalog # FTC3) antibodies, HRP-conjugated anti-mouse IgG (1:4500 dilution; Catalog # HPO5), anti-rabbit IgG (1:3500 dilution; Catalog # HPO3) and anti-goat IgG (1:3500 dilution; Catalog # 105500) antibodies were obtained from Merck India.

### Oligonucleotides

The oligonucleotides were synthesized either by Sigma-Aldrich (USA) or by IDT (USA). The sequences of primers used in qPCR and ChIP experiments are provided in Supplementary Tables 2 and 3.

### ShRNA and overexpression constructs

For the downregulation of *BRG1*, ShRNA constructs were used either against the 3’ UTR or against the coding sequence (catalog # SHCLNG-NM_003072; Sigma-Aldrich, USA). *SMARCAL1* was downregulated using ShRNA construct either against the coding sequence (catalog # SHCLNG-NM_014140; Sigma-Aldrich, USA) or against the 3’ UTR created in our laboratory [37]. SMARCAL1 and BRG1 overexpression constructs were created in our laboratory [36,37]. WIP1 (hwip1 FLAG) and PP2A (pmig-Aalpha WT) plasmid were purchased from Addgene (USA).

### Cell culture and transfection

All cell lines were purchased from National Centre for Cell Science, Pune, India, and cultured as detailed previously [36,38]. Transfections were done using Turbofect (Thermo Fisher Scientific, USA) transfection reagent according to the manufacturer’s protocol. In case of transient transfections, the cells were harvested 36–48 hours post-transfection. For stable transfections, cells were selected in antibiotic containing media 48 hour post-transfection.

### RNA isolation, cDNA preparation and qPCR

Total RNA was extracted and cDNA was prepared using the established protocol [36]. qPCR was performed with 7500 Fast Real-Time PCR system (ABI Biosystems, USA) using gene-specific primers designed for exon-exon junctions using the established protocol [36].

### Immunoblotting

Immunoblotting was done using previously reported protocol [36] and Image J software was used for relative quantitation of protein bands from western blot films.

### Immunofluorescence

Immunofluorescence was performed as explained in Patne et al. [37] and the prepared slides were viewed using confocal microscope (Nikon) under a 60X oil immersion objective.

### Cell cycle synchronization

Cells were synchronized using double thymidine block. Briefly, cells were incubated with 2 mM thymidine for 18 hr. The cells were washed with 1X PBS and cultured in thymidine-free media for 8 hr. Subsequently, cells were incubated for 16 hr in thymidine (2 mM) containing media. The synchronized cells were cultured in fresh DMEM and collected at different times for cell cycle analysis.

### Cell cycle analysis

After washing twice with PBS solution, the cells were fixed with chilled 70% alcohol at −20°C for 24 h. The cell were pelleted by centrifugation (1500 rpm, 3 min), washed twice with PBS solution, incubated with 0.3 mg/ml of RNase A for 30 min at 37°C, and stained with 30 μg/ml PI for 30 min at room temperature. The cell cycle distribution was analyzed using BD FACSCalibur system.

### Chromatin Immunoprecipitation

Chromatin immunoprecipitation was performed using the previously described protocol [36,37]. ChIP-reChIP was done as per the published protocol [39].

### ncRNA complementation experiment

HeLa cells were co-transfected with ShRNA construct targeting the 3’ UTR of *SMARCAL1*/ *BRG1* along with the vector containing the corresponding wild-type or mutated gene. After 36 hr, the cells were treated with 2 μM doxorubicin for 10 min. The cells were harvested and ncRNA was isolated using mirVana^TM^ miRNA Isolation Kit (Thermo Fisher Scientific, USA) as per the manufacturer’s instructions. HeLa cells were treated with doxorubicin and subsequently, washed with 1X PBS 3-4 times. The cells were lysed with 2% Tween-20 in 1X PBS for 20 min at room temperature. After washing with 1X PBS, the cells were treated with 1mg/ml RNase for 15 min at room temperature. The RNase was inactivated by incubating with RNaseOUT (Thermo Fisher Scientific, USA) for 15 min at room temperature. The cells were incubated with 100 ng of ncRNA isolated from transfected cells as per the established protocol [40]. The cells were subsequently processed for immunostaining.

### ATPase assay and CD spectroscopy

ATPase assays were performed using purified ADAAD as published previously [41]. CD spectra was recorded using Chirascan (Applied Photophysics) using the protocol published previously [38]. The CD values were converted into Mean Residue Ellipticity [θ] using the following equation:

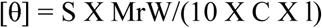

Where S is the CD signal, MrW is the mean residue weight of the DNA, C is the concentration of DNA, and l is the pathlength of the cuvette.

### Statistical analysis

The statistical significance was calculated using Sigma plot. The qPCR data, unless otherwise stated, is presented as average ±s.d of three independent experiments. The western blots were quantitated using Image J software and the data was normalized with respect to β-actin. The quantitated data, unless otherwise stated, is presented as average ±s.d of two independent experiments. In case of imaging experiments, unless otherwise stated, the data is presented as average ±s.d of three independent experiments where n≥100 were counted in each experiment.

## ACKNOWLEDGEMENTS

The authors would like to thank Central Instrumentation Facility, School of Life Sciences for Confocal microscope and FACS. They would also like to thank AIRF, JNU for CD spectrophotometer.

## AUTHOR’S CONTRIBUTION

Conceptualization, R.M. and R.S. Methodology, R.M. and R.S.; Investigation, R.S. K.P., R.R., V.A., D.T.H, T.S, R.K., and I.G.; Writing-original draft, R.M. and R.S.; Writing-review and editing, R.M., R.S., K.P., and R.R..; Funding acquisition, R.M.; Supervision, R.M.

